# Improving Electrochemical Aptasensor Sensitivity for *Bacillus cereus* Spore Detection in Food Safety Applications

**DOI:** 10.1101/2025.08.12.669881

**Authors:** Milica Sentic, Francesco Rizzotto, Zorica Novakovic, Aleksandar Karajic, Brahim Heddi, Jasmina Vidic

**Affiliations:** Université Paris-Saclay, INRAE, AgroParisTech, Micalis Institute, Jouy en Josas, France; University of Belgrade, Institute of Chemistry, Technology and Metallurgy, National Institute of the Republic of Serbia, Belgrade, Serbia; University of Novi Sad, BioSense Institute, 21000 Novi Sad, Serbia; Department of Electrical and Computer Engineering, Tufts University, Medford, MA 02155, USA; Laboratoire de Biologie et de Pharmacologie Appliquée, CNRS UMR8113, École Normale Supérieure Paris-Saclay, Gif sur Yvette, France

## Abstract

Rapid detection of *Bacillus cereus* spores is critical for preventing food contamination and spoilage. Many existing methods detect *B. cereus* cells instead of spores, and cannot be applied directly in foods. Here, combination of aptamers that bind different moieties at the surface of *B. cereus* spore with rapid electrochemical detection is presented. When different DNA aptamers were immobilized on screen printed gold electrode, they showed higher binding capacity for *B. cereus* spores than individual aptamers suggesting a synergic effect. The biosensor exhibited a wide dynamic range (10^2^ - 10^7^ CFU/mL) of low limits of detection (∼1 CFU/mL) using just 15 μL of sample. Validation in spiked salad, using direct spore sensing in rinse water and comparing it to the culturing method, confirmed its sensitivity and specificity. This aptamer-combined approach achieving the rapid (15 min) and single-step detection may also be suitable for detecting other foodborne pathogens.

## 1. Introduction

The World Health Organization European Region estimates that more than 23 million people fall ill from eating contaminated food every year, resulting in 4654 deaths and more than 400,000 disability-adjusted life years (WHO, 2025). However, foodborne diseases are fully preventable, and their burden can be decreased by strengthening prevention, surveillance and management of food safety risks. Besides implementing strict hygienic regulations, the development of innovative pathogen detection methods, that can be applied directly in food, is critical to protect consumers and increase global health security.

Among food contaminating agents, *Bacillus cereus,* a Gram-positive spore-forming bacteria, poses serious problems for both consumers and food industry (Authority, 2019; Jovanovic, Ornelis, Madder, & Rajkovic, 2021). It is the primary microbe associated with baby food, third agent responsible for collective foodborne outbreaks in Europe (Cormontagne, Rigourd, Vidic, Rizzotto, Bille, & Ramarao, 2021; Ramarao, Tran, Marin, & Vidic, 2020) and third leading foodborne pathogen in China (Paudyal et al., 2018). In the United States, 60,000 cases of foodborne diseases caused by *B. cereus* are reported annually (Zhou et al., 2023). *B. cereus,* also known as *Bacillus cereus sensus lato* (s.l.), can cause gastrointestinal diseases ranging from mild to severe, primarily through the production of enterotoxins and emetic toxins that cause diarrhea and emetic syndrome, respectively (Ehling-Schulz, Lereclus, & Koehler, 2019; Ramarao et al., 2020). Moreover, *B. cereus* has been identified as the causing agent of systemic and localized infections, especially in high-risk population, and in rare cases, of toxic shock syndrome affecting the central nervous system (Cormontagne et al., 2021). The most dangerous species of the *B. cereus* group is *Bacillus anthracis*, the etiologic agent of anthrax, which causes a highly lethal disease in both humans and animals (Ehling-Schulz et al., 2019).

*B. cereus*, is widely distributed in the environment and can contaminate any food primarily through soil and air (Ehling-Schulz et al., 2019). In its natural biotopes, *B. cereus* is manly present in form of spores and biofilm (Huang, Flint, & Palmer, 2020). Spores allow bacteria to persist for years or even decades under environmental stressors such as heat, dryness, UV radiation, and chemical disinfectants (Huang et al., 2020; Vidic, Chaix, Manzano, & Heyndrickx, 2020). Spores are frequently detected in vegetation, waters, meat, eggs, milk, and processed food, where they can survive heat-treatment, such as pasteurization, that eliminate other microorganisms. Then, when conditions are favorable, the spores of *B. cereus* germinate and outgrow leading to deterioration of the organoleptic qualities of food or produce toxins that can enter the human or animal body via food.

Official techniques for diagnosing *B. cereus* are culture based (Rizzotto, Khalife, et al., 2023; Vidic et al., 2019). These techniques are often efficient but labor intensive and time consuming. The initial results obtained after 2-3 days indicate the presence of the presumptive *B. cereus* cells, however, confirmation of the specific strain involved may take more than one week (Vidic et al., 2019). Advanced techniques, such as PCR, mass spectrometry, microarray, and genomic sequencing are promising and relatively rapid but require purified genetic material, high technical skills, expensive and intense sample processing (Vidic et al., 2019). Moreover, these diagnostic protocols are not directly applicable to spores, and they require an additional step: induction of spore germination, followed by detection of the resulting vegetative cells (Vidic et al., 2020). Consequently, no culturing or molecular method is available for routine use to screen food for *B. cereus* spores. Nevertheless, various categories of advanced biosensors are currently under development (Fischer et al., 2015; Mazzaracchio et al., 2019; Park, Kong, Lee, Ryu, & Park, 2018; Rizzotto, Marin, Péchoux, Auger, & Vidic, 2023; Vidic et al., 2020; Zhou et al., 2023). Among which, aptamer-based sensing stands-out as one of the most promising strategies that allows for an efficient and specific targeting of whole bacterial spores. This is particularly promising because they are low-cost, easy-to-use and deployable throughout the food manufacturing chain.

Aptamers are short single-stranded nucleic acid sequences capable of selective and specific binding to target, generally, due to their specific 3D structure. They are generated *in vitro* through a process called SELEX (Systematic Evolution of Ligands by Exponential Enrichment) and represent a cheaper and reversible alternative to expensive immuno-assays. Unlike antibodies, aptamers show low immunogenicity, extended shelf-life, and are inexpensive and easy to produce (Léguillier, Heddi, & Vidic, 2024; Manceau et al., 2024). However, despite their attractiveness as recognition elements, aptasensors are still in their early stages and require long-term investigation and optimization for real-life applications (Brown, Brill, Amini, Nurmi, & Li, 2024).

Aptamers selected against *B. cereus* spores can be integrated into diagnostic aptasensors that function without the need for spore germination or lysis. Several aptamers with high affinity and specificity to bind *B. cereus* spores have been selected using cell-SELEX and are available in the literature (Bruno & Carrillo, 2012; Fischer et al., 2015; Zhou et al., 2023). This ability to recognize binding sites on the surface of spores, rather than intracellular biomarkers, should significantly reduce analysis time, representing a major advantage for low-cost and rapid food diagnostics. Although the precise molecular targets of aptamers selected by cell-SELEX remain to be identified and validated, it is unlikely that they bind to the same target, given the organizational complexity of the spore surface and the diversity of the 3D structures of published aptamers.

Therefore, we speculate, here, that using a combination of aptamers that recognize *B. cereus cereus* spores will allow targeting multiple recognition sites at their surface, which will provide an enhance diagnostic sensitivity. For this, we tested three different aptamers that have been previously characterized for their binding to spores of various *B. cereus* strains, including. *B. anthracis*, named BAS6 (Bruno et al., 2012; Mazzaracchio et al., 2019), Apt1 (Fischer et al., 2015) and Apt2 (Fischer et al., 2015). We characterized their structural stability in solution, and demonstrated that they do not hybridize with each other. Binding assays were conducted to compare the signal generated by individual aptamers and aptamer mixtures upon binding to *B. cereus* spores. Then, the mixtures or individual aptamers were employed in an aptasensor for electrochemical detection of spores, and their sensitivity and limit of detection were compared. Finally, the applicability of the aptasensor incorporating the aptamer combination with the best analytical performance was demonstrated through the direct detection of spores in fresh salad samples, requiring minimal sample preparation and providing results within 15 min.

## 2. Materials and methods

### 2.1. Reagents and Materials

All chemicals used in this work were analytical grade. Potassium hexacyanoferrate (III) (ferricyanide), potassium hexacyanoferrate (II) trihydrate, absolute ethanol, sulfuric acid, 6-mercapto-1-hexanol (MCH), glutaraldehyde, potassium chloride (KCl), potassium phosphate (K_3_PO_4_) and Tris (2-carboxyethyl) phosphine hydrochloride (98%, TCEP) were purchased from Sigma-Aldrich (France). 1,1,1,3,3,3-hexamethyldisilazane (HMDS ≥98%) was purchased from Acros Organics (Geel, Belgium). Phosphate buffered saline (PBS), Ph 7.4, was purchased from Lonza (Basel, Switzerland) and used as 1X. Three aptamer sequences were obtained from Eurofins Genomics SAS (Nantes, France). Table 1 summarizes the aptamer sequences used in this study. Their secondary structures were predicted using the mFold web server (Zuker, 2003). All aptamer sequences were purchased as unmodified oligomers, thiolated at 5’end with HO(CH_2_)_6_SS(CH_2_)_6_(T)_5_ group, or modified with TexasRed on 5’ end.

**Table 1.**
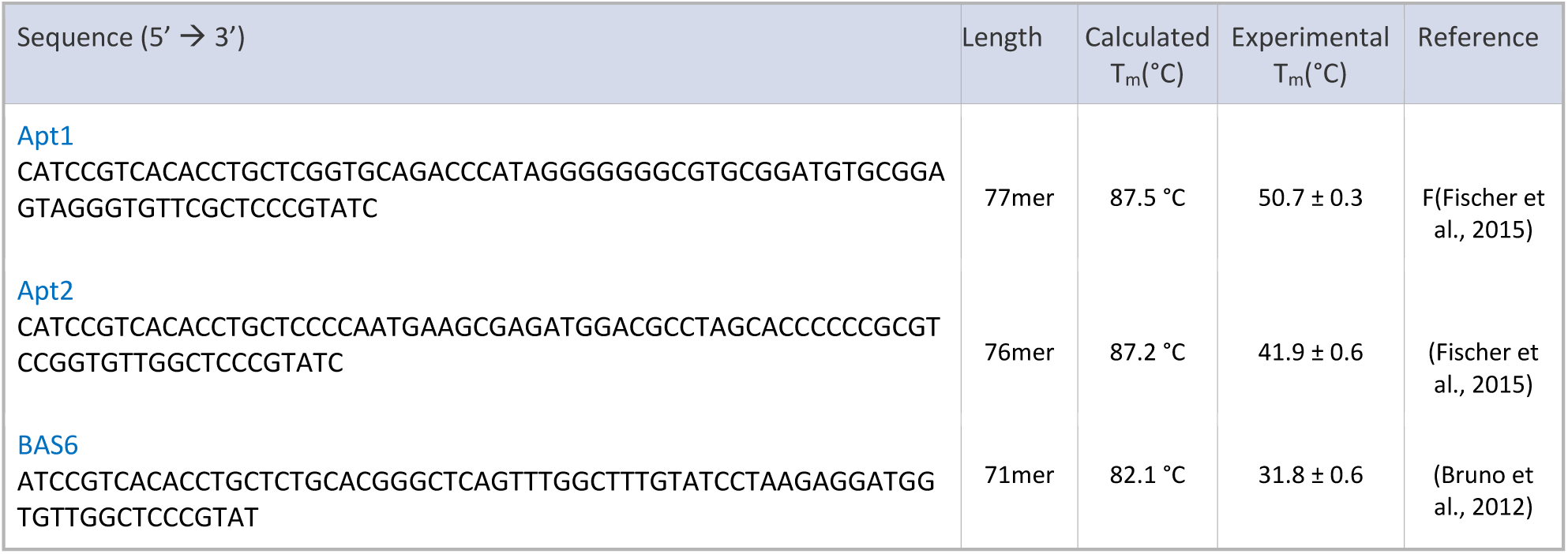
Sequences of ssDNA used in this study.

Upon arrival, the lyophilized pallet of modified aptamer was diluted to a 100 μM stock solution using Milli-Q water and kept at -20 °C until use. Screen Printed Gold Electrodes (GSPE), type BT220, were purchased from Metrohm-DropSens (France). Texas-red aptamer solutions were used at final concentration of 500 nM.

### 2.2. Bacterial stains and growth conditions

Non-pathogenic *B. cereus* strain INRA-SV-S51 isolated from soil was used in this study. *Bacillus subtilis* NDmed strain was used as a control. For routine growth, *B. cereus* was propagated in Luria-Bertani (LB) medium (1% NaCl, 1% tryptone, 0.5% yeast extract) at 37 °C. For sporulation, bacterial cells were cultured in sporulation medium HCT (0.7% casein hydrolysate, 0.5% tryptone, 0.68% KH_2_PO_4_, 0.012% MgSO_4_·7H_2_O, 0.00022% MnSO_4_·4H_2_O, 0.0014% ZnSO_4_·7H_2_O, 0.008% ferric ammonium citrate, 0.018% CaCl_2_·4H_2_O, 0.3% glucose, pH 7.2) (Lecadet, Blondel, & Ribier, 1980) at 30 °C, with rotary agitation set to 200 rpm.

### 2.3. UV Melting experiments

Melting experiments were performed on a Jasco V-750ST UV-visible (UV-Vis) spectrophotometer. Samples of 1-2 µM DNA were prepared by dissolving the stock solution in a mixture of 70 mM potassium chloride and 20 mM potassium phosphate (pH 7.0). For each measurement, the absorbance at 268 nm was recorded as a function of the temperature (15-80 °C). The heating and cooling rate was 0.2°C/min. The determined melting temperature (Tm) represents an average of the result obtained during the heating and cooling cycles.

### 2.4. Circular dichroism experiments

Each aptamer (0.5-1µM) was dissolved in a mixture of 70 mM potassium chloride and 20 mM potassium phosphate (pH 7.0) and circular dichroism (CD) spectra were acquired at 20 °C by using a Jasco J1500 spectropolarimeter. Spectra were recorded from 220 to 320 nm, at 200 nm/min. For each measurement, an average of 3 consecutive scans was taken, the contribution of the buffer was subtracted, and the data were zero-corrected at 320 nm.

### 2.5. Native gel Electrophoresis

The DNA aptamer (1-2 µM) was dissolved in 70 mM potassium chloride-20 mM potassium phosphate (pH 7.0) mixture. Samples were loaded on 20% nondenaturing polyacrylamide gels that was previously spiked with 50% (v/v) glycerol. The gels were visualized by UV-shadowing.

### 2.6. Spectrophotometry

The binding of aptamers to spores of *B. cereus* S51 strain was evaluated using the UV–Vis spectrophotometer Biochrom Libra S22 (Biochrom Ltd., Cambridge, UK). Spores (10^4^ cfu/mL) were mixed with 5 μM of aptamer in 10 mM Tris-HCl, pH 8 and incubated at 4 °C for 4 h with gentle shaking. Then, the spores were centrifuged at 3000 rpm for 5 min, washed three times and resuspended in 10 mM Tris-HCl, pH 8, for analysis. In control experiments, aptamers were replaced with the control oligomer sequence Campy (5’-GGGAGAGGCAGATGGAATTGGTGGTGTAGGGGTAAAA TCCGTAGA-3’).

### 2.7. Construction of electrochemical apta-sensor

The aptasensors were constructed using aptamer sequences modified at the 5’-end with Thiol-C_6_-p(T)_5_. Thiolated-aptamer working solution was prepared by first mixing an aliquot of the 50 µM aptamer solution with the equal volume of 0.5M TCEP. The reaction was proceeded for 1h to ensure the complete reduction of disulfide bonds to thiol. The mixture was then passed through the Spin Desalting Column (7K MWCO, Thermo Scientific, France) and diluted to aptamer concentration of 1 µM with PBS (pH 7.4) for use in functionalization of cleaned gold electrode surface. The SPGEs were functionalized with aptamers according to common procedures previously reported (Watkins, Karajic, Young, White, & Heikenfeld, 2023; Xiao, Lai, & Plaxco, 2007). Prior to functionalization, SPGE surface was cleaned electrochemically by preforming cyclic voltammetry (CV) in 0.05 M H_2_SO_4_ in the potential range from 0 V to +1.1 V at the scan rate of 0.4 V/s (Fig. S1). A total of 50 repetitive scans was performed to ensure the cleanliness of the electrode surface as demonstrated by the overlapping of consecutive cycles. Once the electrochemical cleaning was complete, electrodes were thoroughly rinsed with Milli-Q water and dried. The electrodes were biofunctionalized immediately after the cleaning step by drop-casting 15 µL of the reduced 500 nM thiolated-aptamer solution. The electrodes were incubated for 1h at ambient temperature. The aptamer functionalized SPGEs were rinsed with PBS and incubated overnight at ambient temperature in 5 mM MCH solution prepared in PBS. Finally, functionalized electrodes were copiously rinsed with Milli-Q water prior to electrochemical measurement. Different spore concentrations were incubated during 15 minutes prior to EIS measurement at room temperature. All electrode potentials are given vs screen printed quasi reference Ag/AgCl electrode.

### 2.8. Electrochemical measurements

All electrochemical measurements were performed by a portable PalmSens4 potentiostat/galvanostat/impedance analyzer (PalmSens BV, Netherlands), connected to a laptop equipped with PsTrace 5.11 software. CV measurements were conducted across a potential range from -0.4 to +0.5 V and at a scan rate of 100 mV/s, using PBS solution as a supporting electrolyte and 2.5 mmol/L of [Fe(CN)₆]³⁻^/^⁴⁻ as a redox probe. EIS measurements were recorded in the presence of 2.5 mmol/L [Fe(CN)₆]³⁻^/^⁴⁻ as a redox probe in PBS. EIS was performed by using OCV as the equilibrium potential by applying: 10mV amplitude, number of frequencies: 71, max. frequency: 10,000 Hz, min. frequency: 1 Hz, min. sampling time: 0.5 s, max. equilibrium time: 5 s, OCP max. time was varying from 250 to 500 s for the different sensor layers, stability criterion: 0.001 mV/s. Subsequently, charge transfer resistance (R_ct_) was determined by approximation based on calculating the diameter of the semi-circle of the real part of impedance (Randviir & Banks, 2022). The analytical plot was constructed using signal gain (R (%) =100 × (R_ct_ sample− R_ct_ blank)/R_ct_ blank)) versus a logarithmic concentration range of *B. cereus* spores. The target detection limits (LoD) were calculated as three times the standard deviation of the blank divided with the slope of the obtained calibration curve, while limit of quantification (LoQ) was determined as a 3.3 time the LOD according to the definition by the International Union of Pure and Applied Chemistry (IUPAC) (Long & Winefordner, 1983).

### 2.9. Scanning electron microscopy

Functionalized SPGE carrying thiolated aptamers was incubated with 15 µL *B. cereus* spores water solution (10^4^ CFU/mL) for 15 min at room temperature. Then, electrodes were washed with Milli-Q and mounted on aluminum stubs with carbon adhesive discs (Agar Scientific, Oxford Instruments SAS, Gomez-la-Ville, France). The surface was visualized by field emission gun scanning electron microscopy (SEM FEG) as secondary electrons images (2 keV, spot size 30) under high-vacuum conditions with a Hitachi SU5000 instrument (Milexia, Saint-Aubin, France at facilities located on the MIMA2 platform, INRAE, Jouy-en-Josas, France; https://doi.org/10.15454/1.5572348210007727E12).

Spores in solution were visualized using an Apreo 2C High-Resolution SEM (HRSEM; Thermo Fisher Scientific, Waltham, MA, USA). Samples were prepared following the previously described protocol (Grafskaia et al., 2022). Briefly, *B. cereus* and *B. subtilis* were suspended in PBS, centrifuged at 3000×g for 5 minutes and washed twice with PBS. Fixation was performed overnight at 4 °C using 2.5% glutaraldehyde solution. Following fixation, the samples were centrifuged, and the supernatants were removed. The resulting bacterial pellets were deposited on sterile silicon wafers mounted on 32-mm aluminum stubs with carbon adhesive. Dehydration was achieved through a graded ethanol series (10% to 100% in 10% increments, 5 minutes per step). Subsequently, the samples were incubated for 5 minutes in a 1:1 (v/v) mixture of ethanol and 50% HMDS, followed by incubation in 100% HMDS until complete evaporation. After air-drying, the samples were sputter-coated and imaged under HRSEM at an accelerating voltage of 2 keV and a probe current of 50 pA using an Everhart–Thornley detector (ETD).

### 2.10. Spores labeling using Texas Red-labelled aptamers

Slide-a-Lyzer Dialysis cassettes were purchased from Thermo Scientific (Lissieu, France) (FITC). A volume of 99 µL of *B. cereus* spores (10^7^ CFU/mL) was mixed with 1 µL of Texas Red-labelled aptamer sequences (100 µM) and incubated overnight at 4°C under gentle shaking. After incubation, the samples were washed with PBS and precipitated by centrifugation five times at 3,500 rpm for 3 min. Supernatant was removed by decantation and labeled spores were resuspended in PBS. A control sample containing *B. subtilis* spores at 10^7^ CFU/mL was prepared using the same incubation method and washing steps.

### 2.11. Fluorescent microscopy

Images were acquired with an Axio-Observer Z1 inverted fluorescence microscope (Zeiss) equipped with an AxioCam 807mono digital camera (Zeiss) and fluorescence filters using a ×100 Apochromat objective (Zeiss). Spores stained with TexasRed-aptamers were imaged using the 38 HE filter (excitation: BP 470/40, beam splitter: FT 495, emission: 525/50), and mCherry was imaged using the 45 HE filter (excitation: BP 590/20, beam splitter: FT 605, emission: 620/14). Images were processed using the ZEN software package (Zeiss) and ImageJ.

### 2.12. Spore contamination in Ready-to-Eat salad samples

A sample of the pre-washed salad (∼50 g) was placed in a sterile Becher and inoculated with 1 mL of *B. cereus* spores water suspension (∼10^3^-10^5^ CFU/mL). The salad was gently mixed and then flooded in 250 mL commercial mineral water for 15 min at 4°C to simulate spore transfer. The transfer was performed at refrigerated temperature to mimic supermarket storage conditions and to prevent possible germination. Afterword, the salad was removed, and the spore containing water was collected and centrifugated. The pallet was resuspended in 1 mL fresh mineral water. The protocol aligns with established microbiological protocols for *B. cereus* food contamination modeling (Bacteriological Analytical Manual, BAM)(Tallent, Knolhoff, Rhodehamel, Harmon, & Bennett, 1998). Serial dilutions of the resulting spore-containing samples were analyzed using the aptamer-based biosensors. In parallel, spore enumeration were performed via plate counting to enable evaluation of biosensor performance. In control experiments, *B. cereus* spores were replaced with spores of *B. subtilis*.

## 3. Results and Discussion

### 3.1. Aptamer characterization

The originality of this work lies in the development of electrochemical aptasensors based on a mix of aptamers targeting different binding sites on the surface of the *B. cereus* spores in a non-competitive manner. The objective was to improve the electrochemical signal generated upon aptamer binding to captured spores by introducing multiple oligonucleotide sequences, as compared to the traditional single-sequence approach. A gel migration assay showed that Apt1, Apt2, and BAS6R aptamers (Table 1) do not bind to each other (Fig. 1a), thereby eliminating concerns of their spontaneous hybridization that could compromise target recognition when these sequences are used in a mixture. In dilute solution (water solution containing 70 mM KCl), CD spectra of the three aptamer sequences show a characteristic positive peak at 275 nm and a negative peak at 240 nm for folded DNA molecules (Fig. 1b) The folding was confirmed by 2D modeling of the three aptamers using the mfold software, which generated several possible structures for Apt1 and Apt2 and one for BAS6 (Fig. S2). The variability in conformations and the loop sites of the three aptamers suggests that they can bind at different sites on the spore surface. However, the existence of multiple 2D structures for Apt1 and Apt2 can be a drawback for their applications, as only one conformation may possess the necessary recognition capability. In addition, UV melting experiments demonstrated that Apt1 and Apt2 were more stable then BAS6 which can be partially unfolded at the room temperature (Table 1). Considering that aptamer conformation is essential for target binding, the low thermal stability of BAS6 may compromise the accuracy and reproducibility of the sensors.

**Figure 1.**
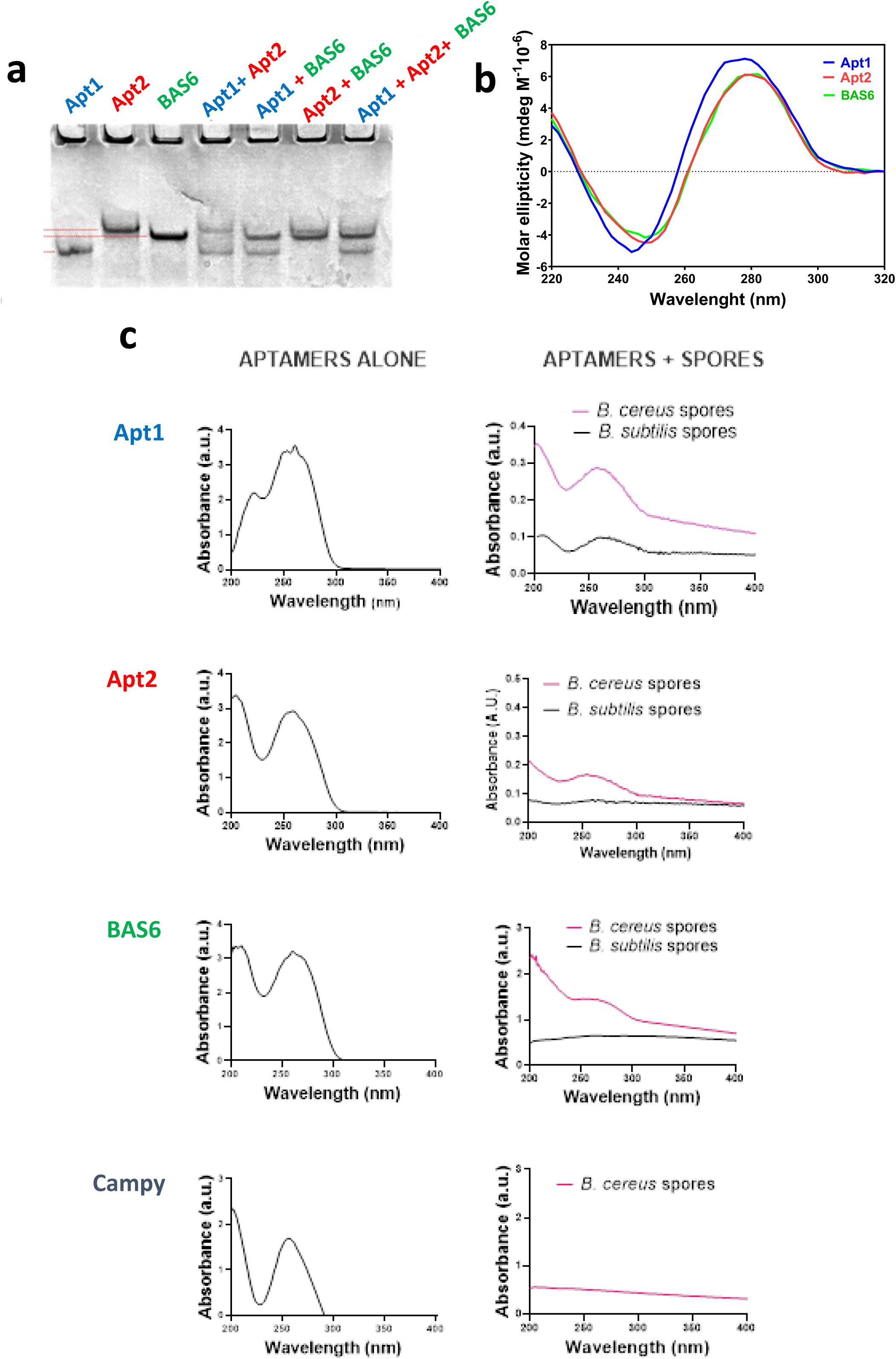
(**a**) Native gel electrophoresis of different aptamer sequences and their mixtures. The name of the sequences loaded in the gels are listed on top of each well. (**b**) CD spectra of Apt1 (blue), Apt2 (red) and BAS6 (green). (**c**) Spectrophotometric detection of BAS6, Apt1 and Apt2 aptamer alone (left panel) and bound to spores of *B. cereus* or *B. subtilis* (right panel). Campy oligonucleotide was used as a negative control. Binding experiments were performed with 10^4^ CFU/mL of spores and 10µM aptamer.

To confirm that the aptamers Apt1, Apt2, and BAS6 bind to spores of the *Bacillus cereus* strain, the spores were incubated with each aptamer and analyzed by UV–Vis spectroscopy. As nucleic acids, aptamers exhibit strong UV absorbance around 260 nm, in contrast to spores, which do not absorb significantly at this wavelength (Fig. 1c). However, when *B. cereus* spores were incubated with aptamers and thoroughly washed, a peak at 260 nm was observed, indicating aptamer binding (Fig. 1c). In contrast, no peak around 260 nm was detected when aptamers were incubated with unrelated spores of *B. subtilis*, suggesting no binding to the control and thus confirming the specificity of the interaction. The specificity was confirmed by incubating *B. cereus* spores with the control oligonucleotide Campy, as no binding was observed in this case either (Fig. 1c). This set of experiments was performed in Tris buffer (pH 8), while Apt1 and Apt2 were selected in a sodium acetate buffer, pH 5 (Fischer et al., 2015), and BAS6 in a Tris buffer containing NaCl and MgCl_2_ salt, pH 7.5 (Bruno et al., 2012). It appears, therefore, that aptamers’ functional structures remain active across a range of pH and salt concentrations.

### 3.2. Verification of aptamers’ synergistic effect

To evaluate whether combining two or three aptamers yields a synergic effect and enhances affinity for *B. cereus* spore binding, the aptamers molecules modified with the fluorescent reporter Texas Red were used for spore staining (Fig. 2a). After removal of unbound aptamers, the thoroughly washed spores were visualized by epifluorescence microscopy, and their staining intensity was analyzed using ImageJ. Ultrastructural visualization of *B. cereus* S51 spores showed their distinct morphological characteristics such as ellipsoidal shape, typical length of 2-3 µm long, with a rugged coat and a prominent exosporium (Figs. 2b and S3). The obtained fluorescent imaging results suggest that selected oligonucleotide sequences can bind both, the core and the exosporium of *B. cereus* spores (Figs. 2c and 2d). Particularly BAS6, alone or in a mixture, marked exosporium which is in line with our previous report on BAS6 interaction with *B. cytotoxicus* spores (Rizzotto, Marin, et al., 2023). The binding of aptamers to exosporium was confirmed by quantification by image analysis of the respective proportion of fluorescence in the stained spores confirmed (Fig. S4). The total molar concentration of each individual aptamer or their combinations (double or triple) was kept constant at 1 µM in order to compare fluorescence intensities. As shown in Figs. 2e, the lowest fluorescence was observed with Apt2 while other aptamers alone or in mixtures showed similar mean of fluorescence intensity over the whole spore area. Accordingly, no synergistic effect, but rather a cumulative one, was observed when multiple aptamers bound to *B. cereus* spores. We next sought to verify whether aptamers can show synergistic effects when immobilized on the electrode surface.

**Figure 2.**
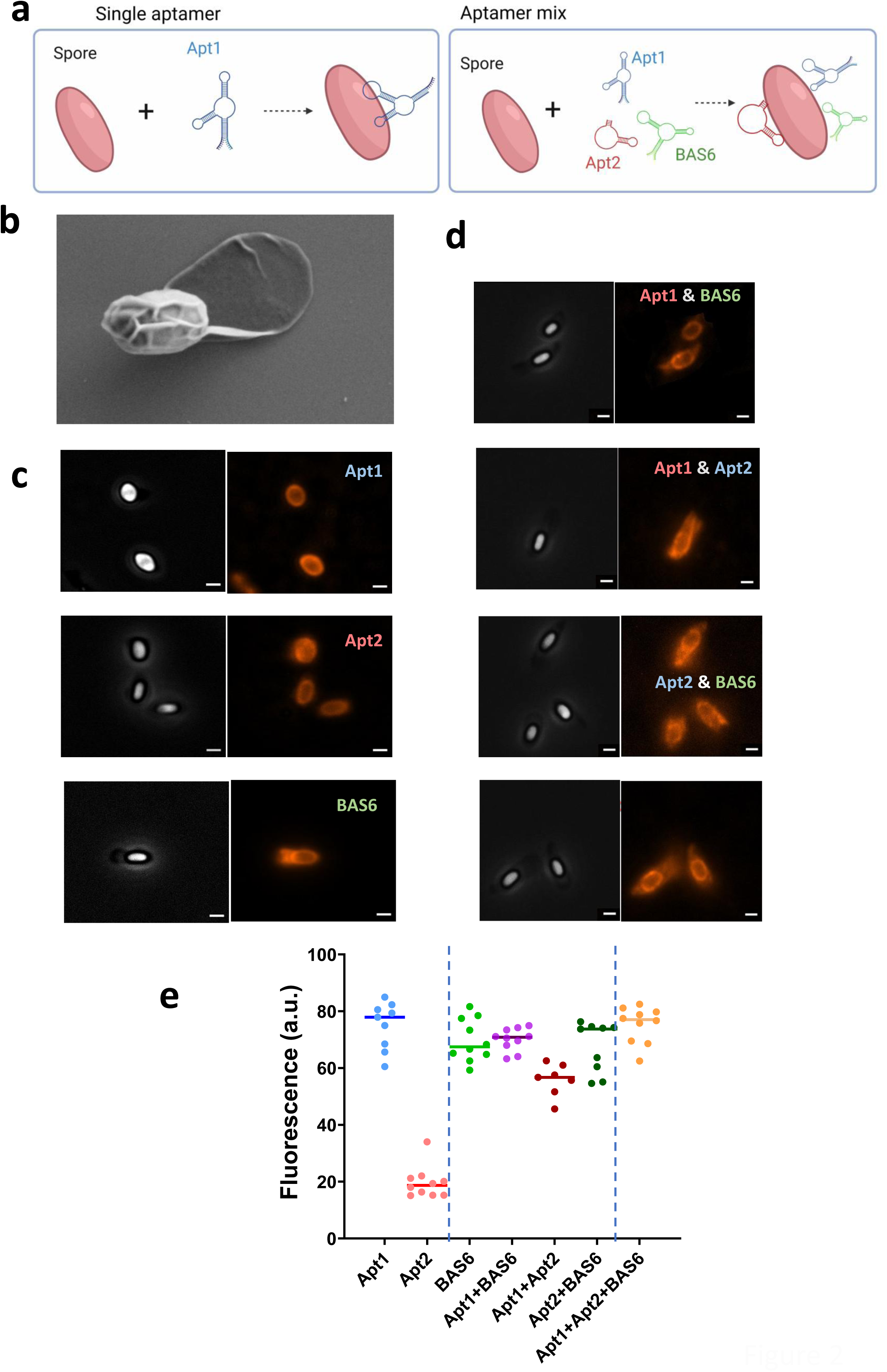
(**a**) Schematic representation of the concept of signal enhancement by using aptamer mixtures for spore detection. (**b**) SEM visualization of B. cereus S51 spore. Scale bar, 1µm. (**c**) Representative fluorescence images of *B. cereus* S51 spores stained with individual aptamers modified with Texas Red at 5’. Scale bar, 1 µm. Left panels show bright field images of spores. (**d**) Representative fluorescence images of *B. cereus* S51 spores stained with combinations (double or triple) of aptamers modified with Texas Red at 5’. Left panels show bright field images of spores. Scale bar, 1 µm. (**e**) Scatter plot of the fluorescence intensity/spore from at least 10 spores as in (**c, d**) determined by ImageJ software. The bar represents the median.

### 3.3. Fabrication and characterization of aptamer-modified electrodes

Fig. 3a illustrates the two-step fabrication process of the aptamer-modified SPGE and the spore capturing. First, the surface of the electrochemically cleaned SPGE was functionalized with the 5’-alcylthiolated-aptamer by using the drop-casting method. This step allowed formation of a self-assembled monolayer of aptamer molecules on gold substrate. Previous works showed that long-chain alkylthiolates can be stably immobilized on the gold surface due to the lateral van der Waals interaction between neighboring monolayer molecules (Bain, Troughton, Tao, Evall, Whitesides, & Nuzzo, 1989; Watkins et al., 2023). In the second step, MCH was employed as a passivation molecule to block unoccupied areas of the gold surface following aptamer binding and reduce the nonspecific adsorption. In addition, MCH was used as the spacer between neighboring aptamer molecules to provide optimal aptamer-aptamer distance for sterically unimpeded binding event with the target spore (Watkins et al., 2023). The SPGE functionalization was characterized by CV using the ferro/ferricyanide as the redox couple in PBS as the supporting electrolyte. Fig. S5 shows a gradual decrease in redox-peak current intensity and an increase in the peak-to-peak separation following electrode incubation with aptamers and MCH. This indicates an increase in the charge transfer resistance of the ferro/ferricyanide species, due to the formation of the mixed monolayer. SEM visualization of the functionalized working electrode suggests the presence of an organic layer over the gold surface, with some defects due to the inherent roughness of the gold (Fig. 3b and Fig. S6). Indeed, the BSE mode for Z-contrast shows different signal intensities on the working electrode surface (Fig. S6), indicating presence of organic matter which appear darker (low Z). Finally, to verify whether the attached aptamers retain their recognition capability, the functionalized electrode was incubated with *B. cereus* spores for 15 minutes, thoroughly washed, and then examined by SEM. Fig. 3c and Fig. S6 shows that spores were indeed captured on the functionalized surface, indicating that the immobilized aptamer kept its functional conformation after immobilization.

**Figure 3.**
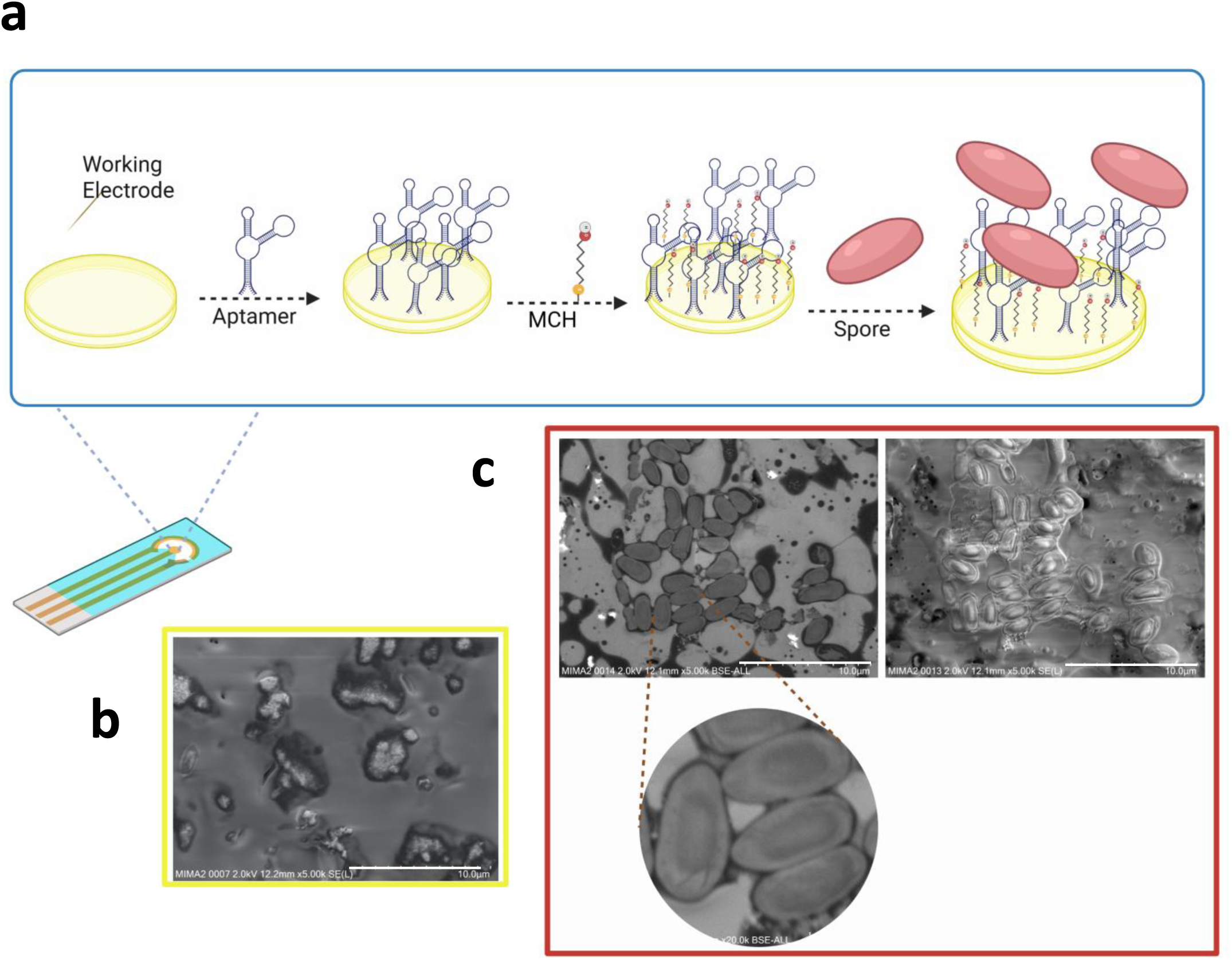
(**a**) Schematic concept of SPGE functionalization by using aptamers and MCH, and subsequential spore sensing. (**b**) SEM image of the electrode functionalized with aptamer BAS6 and MCH. Scale bar, 10 µm. (**c**) SEM images of spores captures on the electrode surface functionalized with BAS6 aptamer and MCH. Scale bar, 10µm.

### 3.4. Electrochemical detection of spores

The comparative detection test was performed with SPGEs functionalized with single or multiple aptamers (at constant total concentration of 1 µM) and exposed to 10^5^ CFU/mL of *B. cereus* spores for 15 min. EIS was used to characterize the spore binding in a solution of the 5mM ferro/ferry cyanide in PBS. Fig. 4a shows the degree of signal gain for single, double or triple aptamers calculated for charge transfer resistance R_ct_. R_ct_ reflects modification of the electrode surface due to target capturing by the molecules of aptamers on the electrode surface since it depends on the dielectric and insulating properties at the electrode/electrolyte interface. A clear synergic effect was observed with electrodes carrying double Apt1+BAS6 and triple Apt1+BAS6+Apt2 aptamers that provided a significant enhancement of the signal gain compared to single aptamers or doubles involving Apt2.

**Figure 4.**
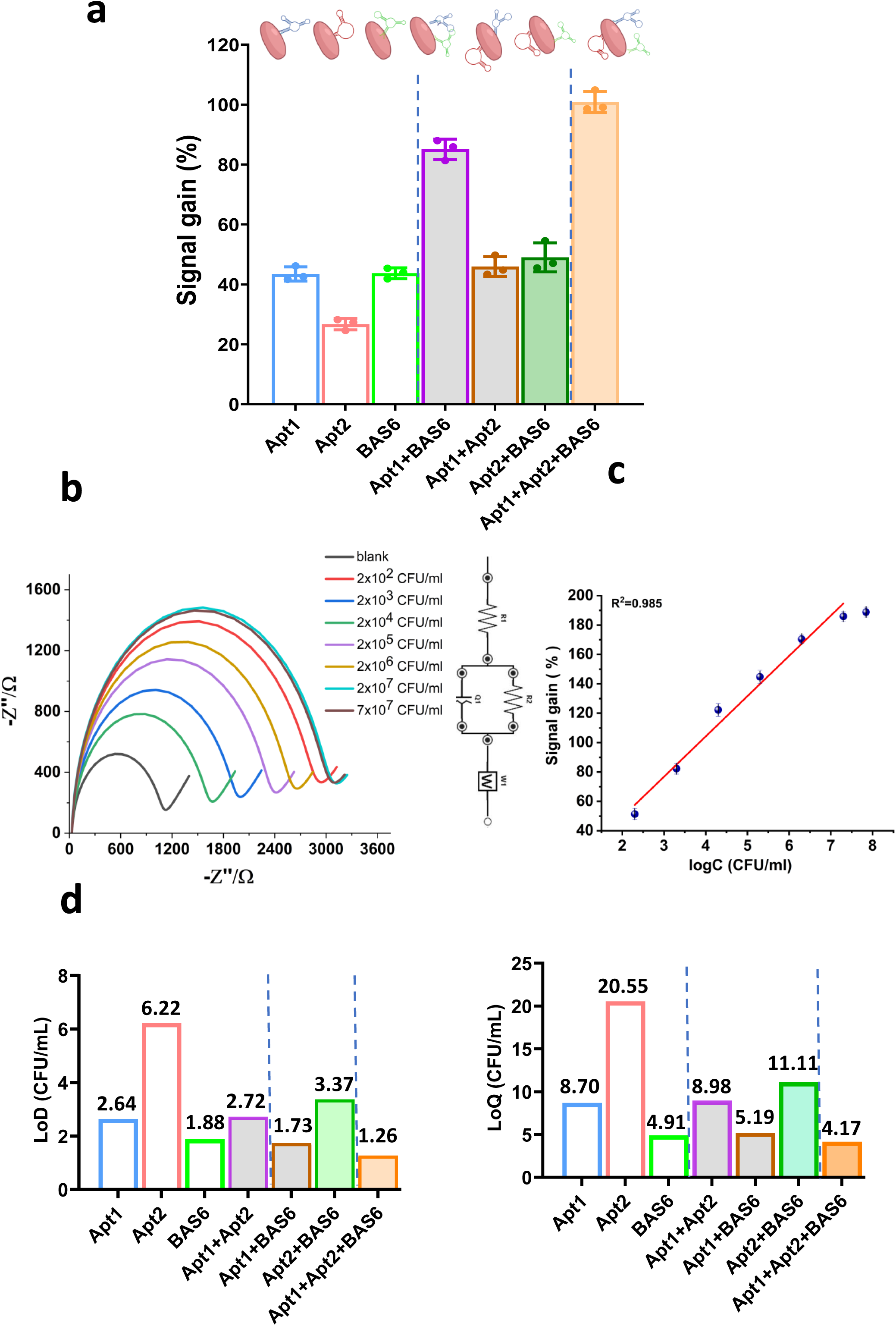
(**a**) Signal gain calculated as 100 × ((R_ct_ sample− R_ct_ blank)/R_ct_ blank) of *B. cereus* spore obtained by the electrochemical biosensor based on single, double or triple aptamers. (**b**) Nyquist plots of impedance spectra taken in PBS with ferro/ferri cyanide redox couple in PBS with various concentrations of *B. cereus* spores using the triple-aptamer biosensor. Insert, the equivalent circuit used for fitting impedance spectra. (**c**) Corresponding calibration curves obtained by plotting the signal gain in a function of *B. cereus* spore concentrations. Data points are the mean values obtained in 3 independent experiments ± SD. (**d**) LoD and LoQ of *B. cereus* spore detection by single or multiple aptamers immobilized on a SPGE.

Figure 4b shows a serios of Niquis plots recorded in a solution containing an increasing concentration of *B. cereus* spores. The obtained data were fitted using an equivalent circuit shown in the inset of Fig. 4b and the corresponding R_ct_ values were calculated. A linear increase in Rct value was observed with increasing the spore concentration from 10^2^ to 10^7^ CFU/mL. As expected, the best analytical properties were obtained with SPGE functionalized with the triple aptamers that exhibited increase of signal gain across all individual aptamers. The triple-aptamer biosensor showed (*R^2^*) of 98.5% and the regression equation *y* = 4.075 *x* + 2.24, where *x* is log (spore concentration in CFU/mL), *y* is signal gain, % (Fig. 4c). The calculated limit of detection (LoD) was 1.26 CFU/mL while the limit of quantification (LoQ) was 4.16 CFU/mL when detection was performed with three-aptamer electrodes. LoD and LoQ of other sensors are presented in Fig. 4d. The triple-aptamer used in the electrochemical aptasensor exhibited, thus, the highest efficiency and was further employed for spores’ detection in a food sample.

### 3.5. Detection of *B. cereus* spores in Ready-to-Eat salad samples

To ensure the analytical reliability of the tripe-aptamer electrochemical sensor, its performance was validated through a study of spiked green salad. To mimic a real-world contamination scenario, ready-to-eat salad samples were inoculated with known concentrations of *B. cereus* spores (10^1^ CFU/mL - 10^7^ CFU/mL), soaked in commercial mineral water to simulate the washing step and the obtained effluent wash-water was analyzed by the triple-aptamer based biosensor (Fig. 5a and Fig. S7). The measurements were performed in solution of the ferro/ferry cyanide redox couple that has been prepared in mineral water. The corresponding Nyquist plots of the impedance spectra are reported in Fig. 5b. The controls performed with *B. subtilis* spores showed significantly smaller variations in Nyquist plots (Fig. 5c). The fitting of the spectra was satisfactory to the same level of that reported in Fig. 4b for PBS as a supporting electrolyte. Fig. 5d shows the calibration curve obtained from the quantification of the signal gain for *B. cereus* spores for each spore concentration. The obtained R_ct_ values present the average of the three independent sensors with the relative standard deviation of ca. 4%. Signal gains plotted against the logarithm of target spore concentrations was linear with a correlation coefficient (*R^2^*) of 99% and the following regression equation: y = 5.46 x + 1.70. For comparison, the insignificant variation of the signal gain was obtained when the triple-aptamer sensor was tested with the same range of concentrations of *B. subtilis* spores used as a negative control (Fig. 5d), confirming the sensor high selectivity towards *B. cereus* spores. Interestingly, due to the better conductivity of the mineral water than that of PBS, the sensitivity of detection in rinse water was exceptional with LoD of 0.9373 CFU/mL and LoQ of 3.0933 CFU/mL. Together, the specificity and selectivity of *B. cereus* spore detection from inoculated salads strongly suggests the reliability of the triple-aptamer biosensor for food safety screening.

**Figure 5.**
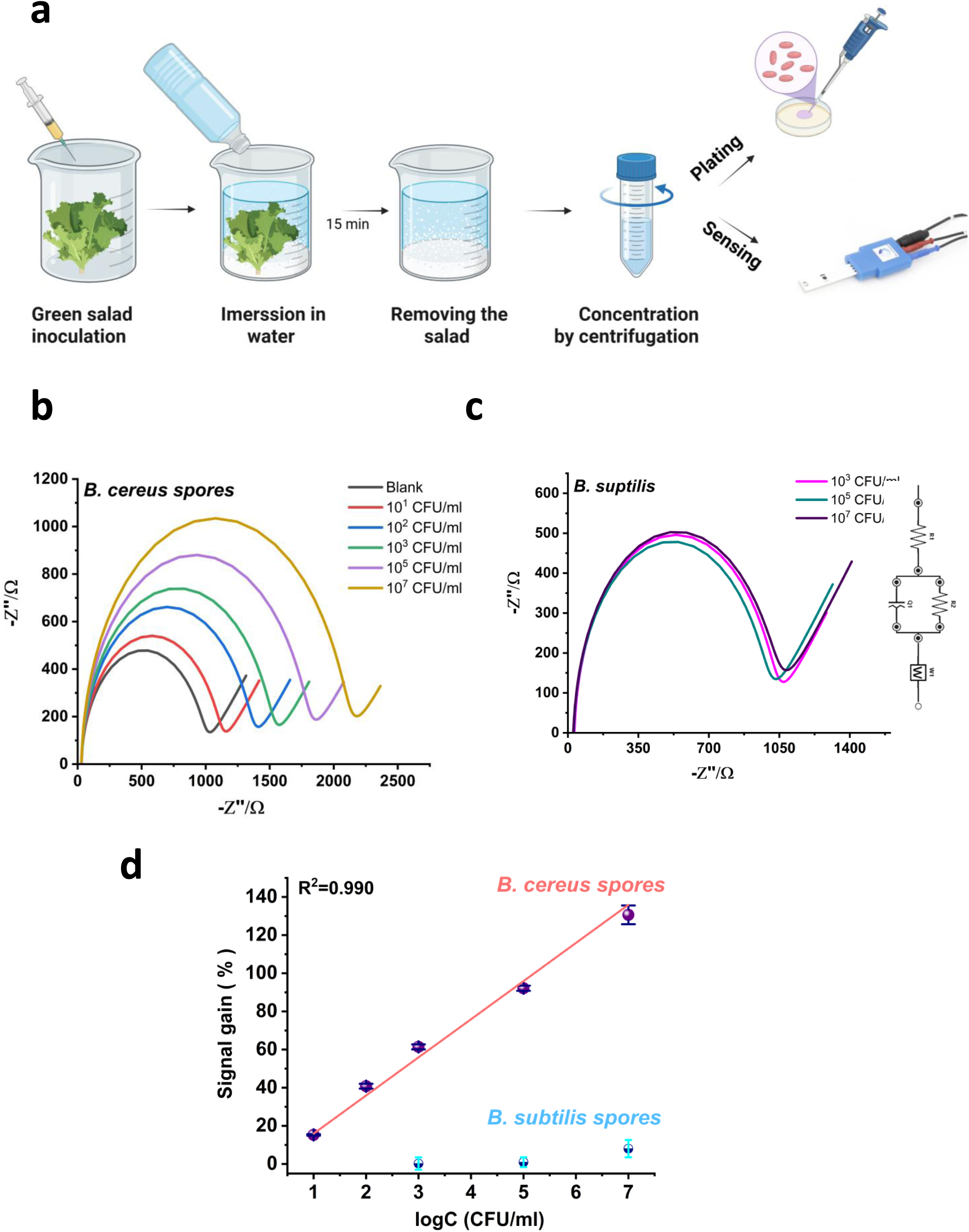
(**a**) Schematic representation of the triple-aptamer based assay building protocol applied to detect *B. cereus* spore in contaminated ready-to-eat salad samples. To mime a real-world contamination scenario, ready-to-eat salad samples were inoculated with known concentrations of *B. cereus* spores, soaked in commercial mineral water to enable transfer and tested by the aptasensor. Results were verified with plating experiments. Nyquist plots of impedance spectra taken with ferro/ferri cyanide redox couple in mineral water with various concentrations of *B. cereus* spores (**b**) or *B. subtilis* spores (**c**) using the triple-aptamer biosensor. Insert, the equivalent circuit used for fitting impedance spectra. (**d**) Calibration curves were obtained by plotting the signal gain in a function of *B. cereus* and *B. subtilis* spore concentrations. Data points are the mean values obtained in 3 independent experiments ± SD.

## 4. Conclusion

The combination of three aptamers and an impedance-based detection strategy was presented in this study as a novel approach for the diagnostic of *B. cereus* spores directly from food samples, without the need for bacterial enrichment, culturing, or biomarker purification. The use of an aptamer mixture effectively increased the sensitivity of electrochemical spore detection compared to single aptamer. This finding suggests that lateral interactions between aptamers within the monolayer formed on the gold electrode surface may play an important role in aptasensor stability and efficiency. The method enabled rapid and ultra-sensitive detection of *B. cereus* spores at concentrations as low as 1-2 CFU/mL within 15 minutes. This triple-aptamer biosensor has the potential to be used by the food industry and regulatory agencies to monitor food quality, especially considering that different *B. cereus* strains can cause food spoilage, gastrointestinal diseases, and infections. The results support the use of aptamer mixture for detection of targets exposing multiple epitopes such as bacteria or eukaryotic cells.

## Supporting information

Supplementary fugures

## CRediT authorship contribution statement

Milica Sentic: Investigation, Formal analysis, Visualization, Writing – review & editing. Francesco Rizzotto: Investigation, Formal analysis, Writing – review & editing. Zorica Novakovic: Formal analysis, Writing – review & editing. Aleksandar Karajic: Writing – review & editing. Brahim Heddi: Investigation, Formal analysis, Resources, Writing – review & editing. Jasmina Vidic: Conceptualization, Funding acquisition, Writing – original draft, Writing – review & editing.

## Acknowledgments

This work was supported in part by the French National Agency for Research (ANR-21-CE21-0009 Siena projects) and the European Union (grant agreement no. 101135402, Mobiles project and grant agreement no. 872662, IPANEMA project). M.S. acknowledges the support of the Ministry of Science, Technological Development and Innovation of the Republic of Serbia, Contract No: 451-03-136/2025-03/200026. We thank Thierry Meylheuc for its expertise and the MIMA2 platform for access to electron microscopy equipment (MIMA2, INRAE, 2018. Microscopy and Imaging Facility for Microbes, Animals and Foods, https://doi.org/10.15454/1.5572348210007727E12).

