## Supplementary fugures for "Improving Electrochemical Aptasensor Sensitivity for *Bacillus cereus* Spore Detection in Food Safety Applications"

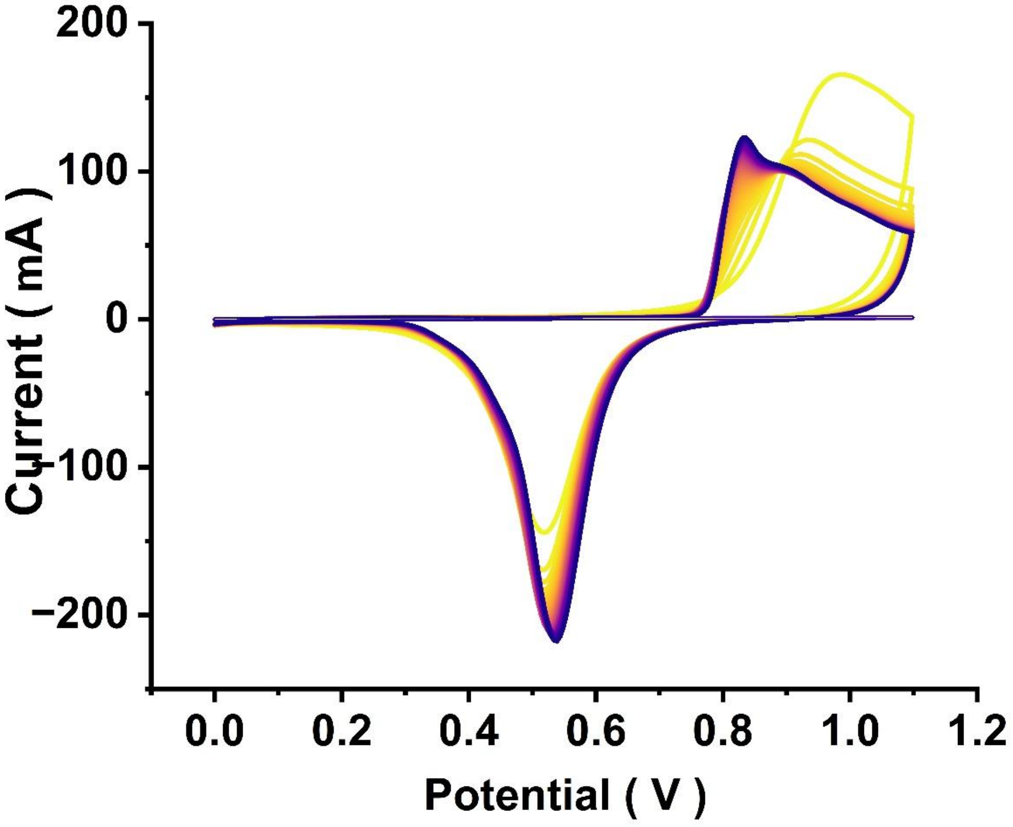


**Figure S1.** SPGE electrode cleaning with 0.05 M $H_2SO_4$ at the scan rate of 400 mV/s and potential window from 0 V to 1.1 V.

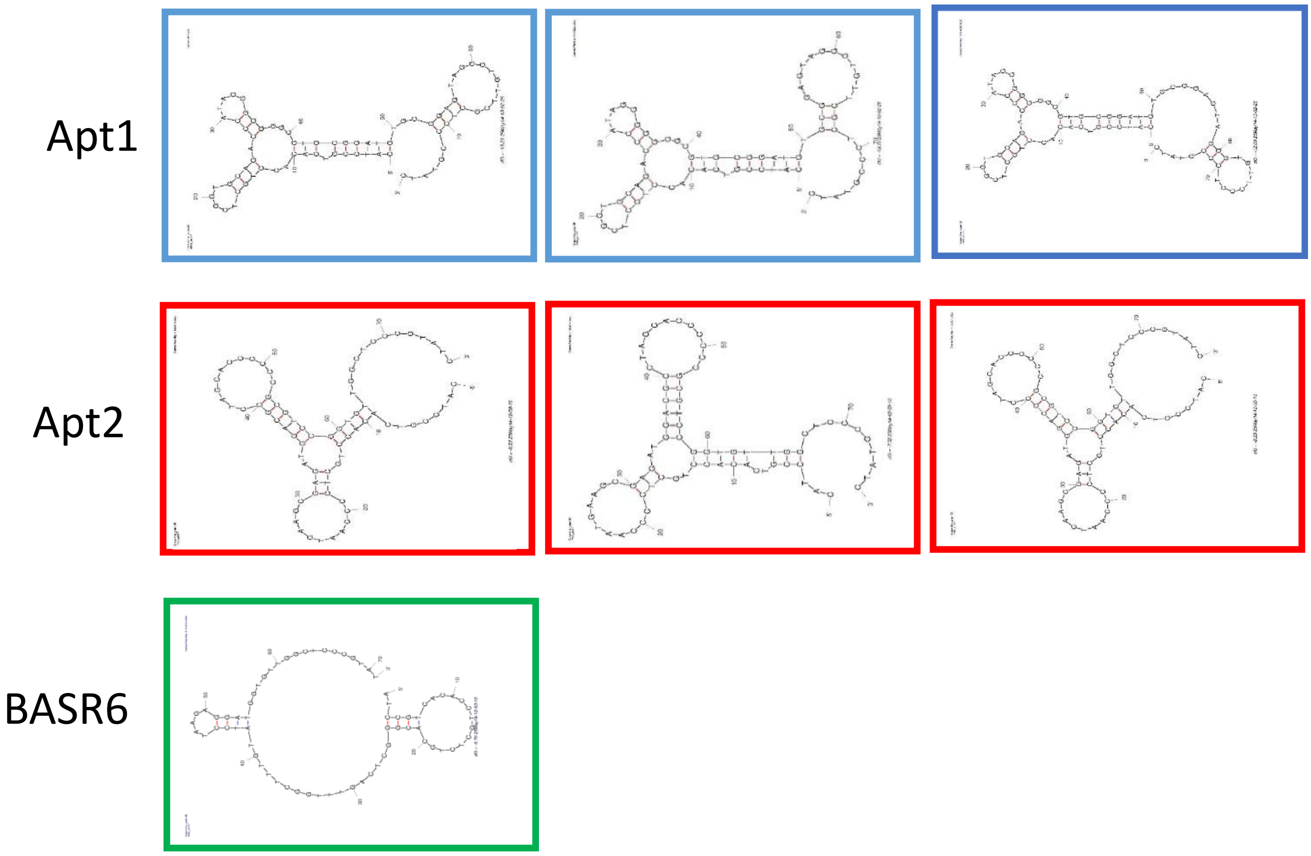


**Figure S2.** Folded structures for the Apt1, Apt2 and BAS6 aptamers as modelized by the mfold software.

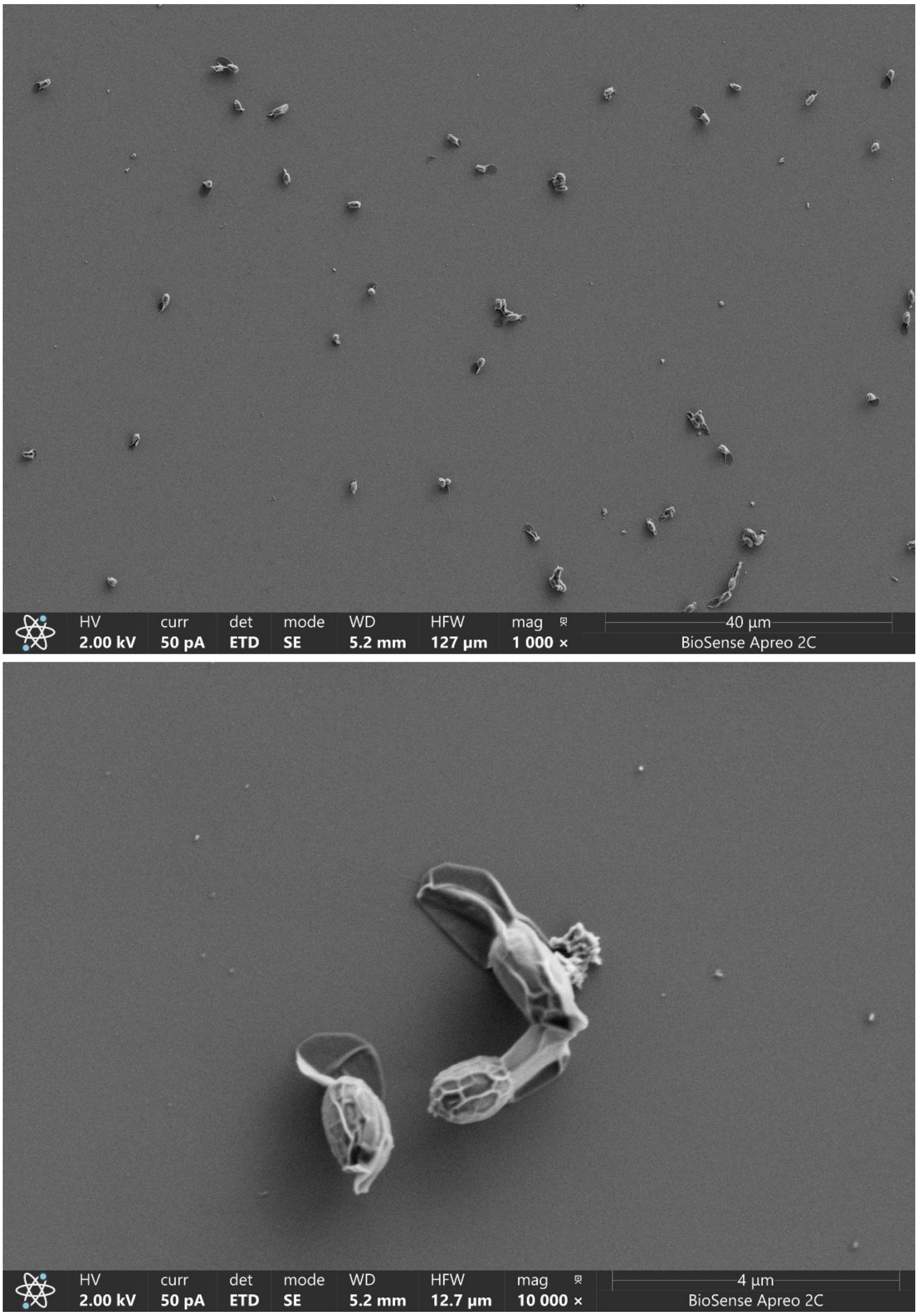


**Figure S3.** Scanning electron microscopy of *B. cereus* S51 spores.

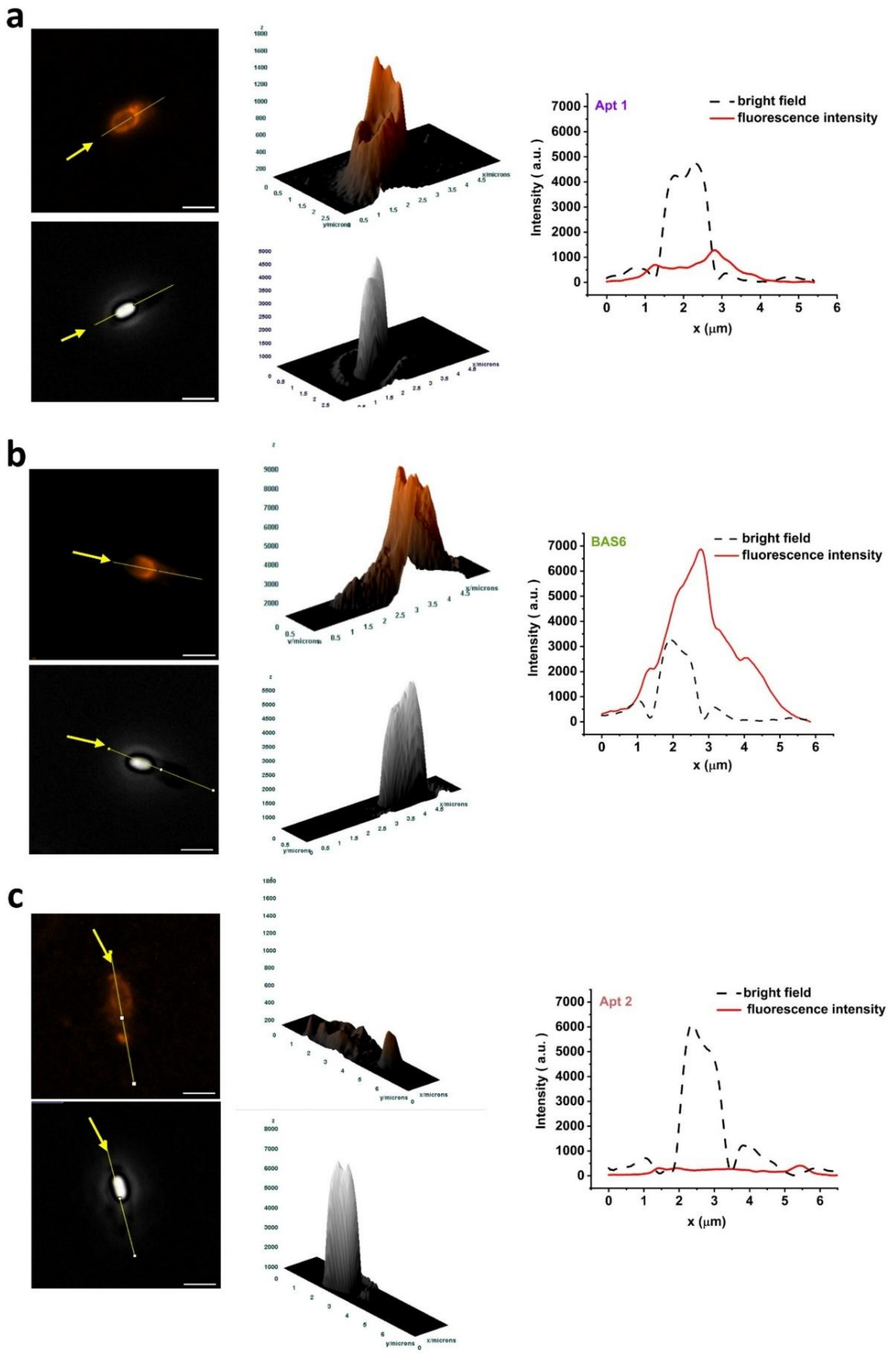


**Figure S4.** Left, representative fluorescence images of *B. cereus* spores stained with modified aptamers at 5' with Texas Red and bright filed images of *B. cereus* spores, scale bar 2 µm. Middle 3D surface plots for yellow lines drown on the left images. Right, corresponding intensity profiles for the same lines drawn on the left images.

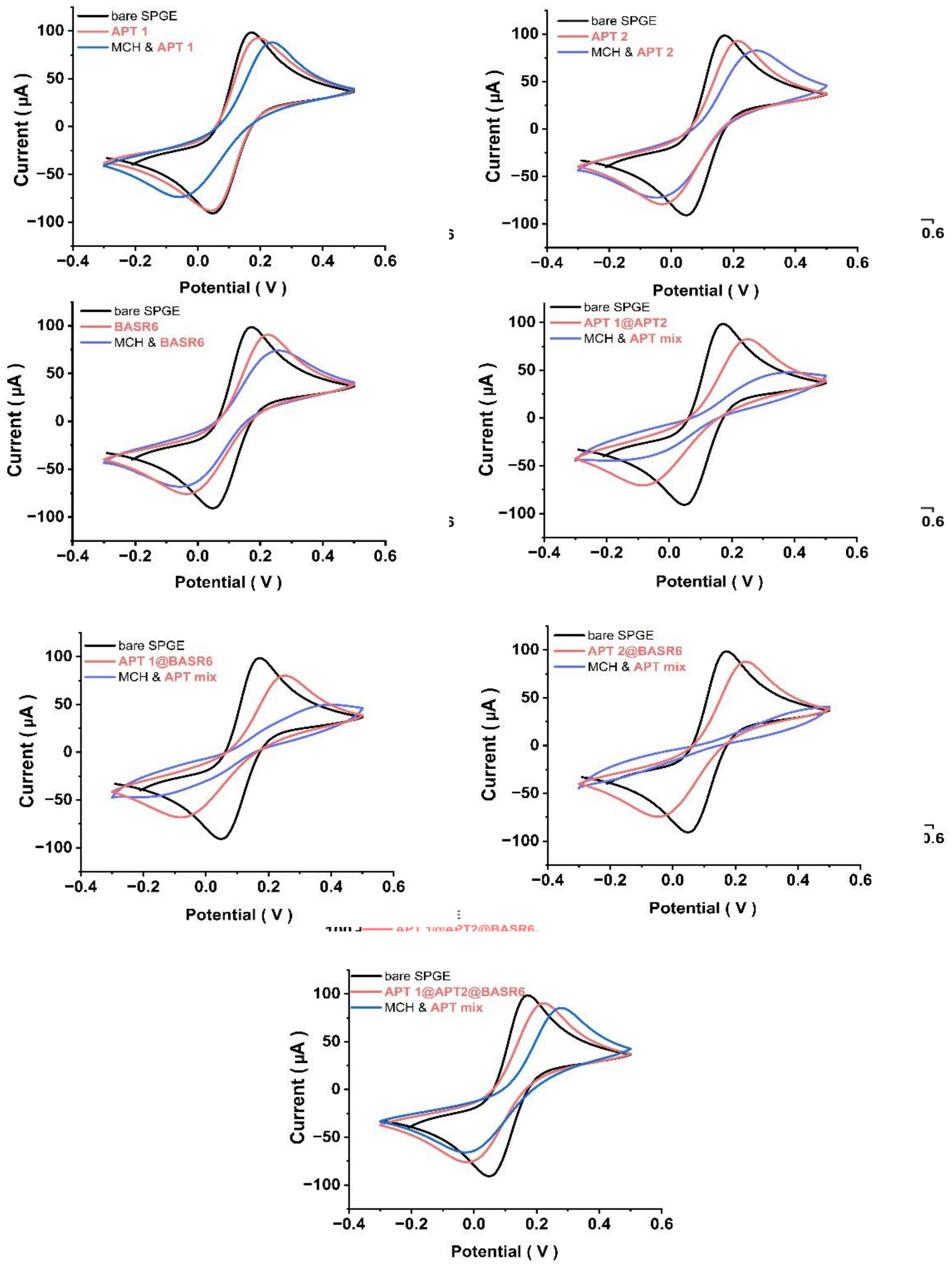


**Figure S5.** Cyclic Voltammograms recorded in 5 mM ferro/ferricyanide in PBS using SPGEs before their functionalization (bare), after immobilization of the aptamers and with aptamer sequence with MCH monolayer at the scan rate of 100 mV/s and potential window from -0.3 to 0.5 V. .

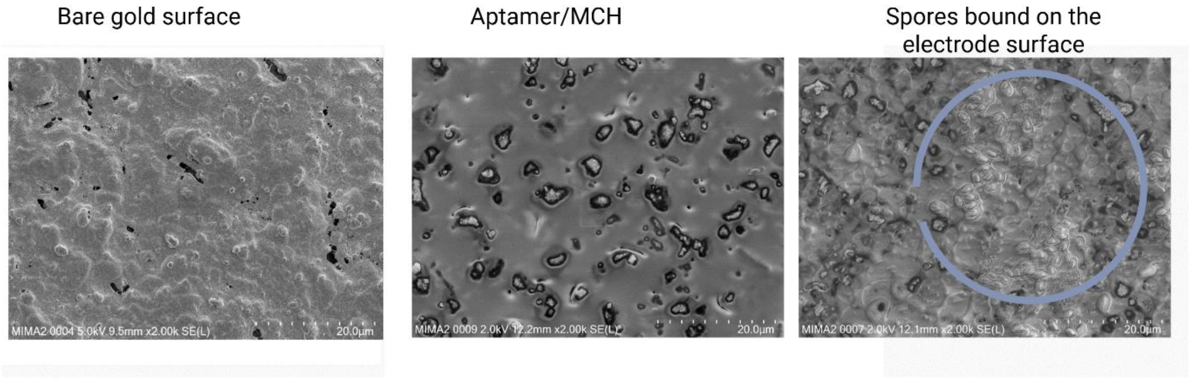


**Figure S6.** SEM images of SPGE: before (Bare gold surface) and after subsequential incubation with 1 µM aptamer BAS6 and 5 mM MCH (Aptamer/MCH). Finally, the modified electrode upon 15 min incubation with $10^5$ CFU/mL of *B. cereus* spores efficiently recognized and captured them. The circle contains many spores (right panel).

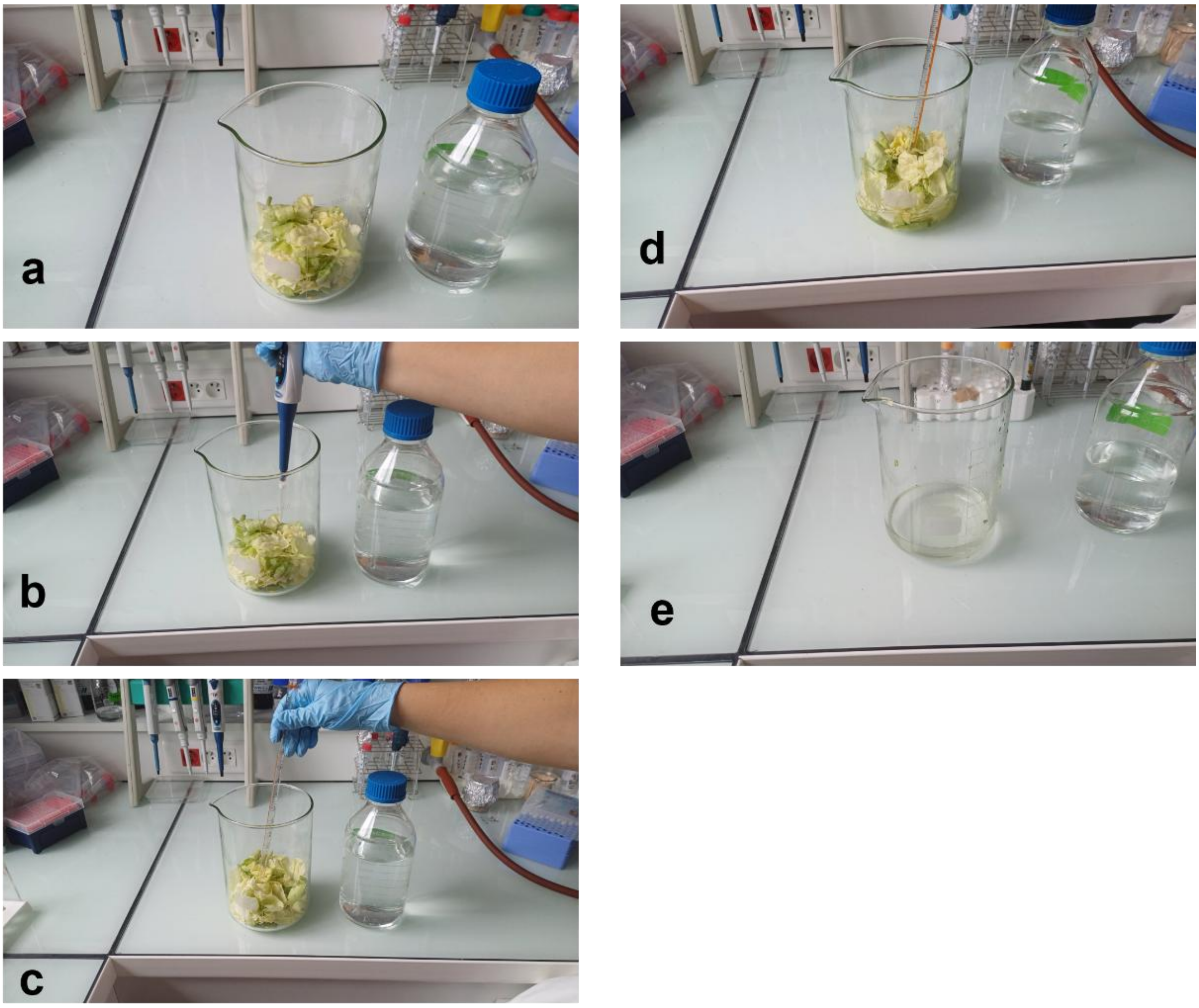


**Figure S7.** To inoculate Ready-to-Eat salad samples, 50 g of pre-washed salad leaves was placed in a sterile Becher (**a**) and 1 mL of *B. cereus* spores water suspension was added (**b**). The salad was gently mixed (**c**) and then flooded in Volvic for 15 min to simulate spore transfer (**d**). The salad was removed (**e**) and the water containing spores was collected and centrifugated.
